# Oblique Thick Light-Sheet Microscopy with Spatiotemporal 3D Localization for RNA Imaging in Intact 50 μm-Thick Sections

**DOI:** 10.1101/2025.06.09.658547

**Authors:** Lin Cai, Qihang Song, Dongjian Cao, Qihuan Li, Wancheng Wu, Yinghao Xiao, Xingguang Chen, Shujing Huang, Jie Yang, Yingjun Zhang, Zhen-Li Huang

## Abstract

Neuron typing in intact brain sections demands 3D imaging reconciling thickness compatibility with large-area coverage—a persistent challenge for RNA imaging methods limited by either restricted fields-of-view (<1 mm^2^) or thin-section constraints (<20 μm). Here we introduce oblique thick light-sheet microscopy integrated with spatiotemporal 3D localization, enabling multiplexed RNA imaging across 50 μm-thick tissue sections and centimeter-scale regions. The 45° illumination geometry achieves optical-interference-free imaging in >100 μm-thick tissues during single-plane scans, while a spatiotemporal localization algorithm recovers submicron resolution by pinpointing fluorescence in situ hybridization (FISH) spots within volumetric excitation. This strategy reveals layer-spanning neuronal distributions and region-specific expression gradients unattainable with conventional RNA imaging approaches. By resolving the historical trade-off between imaging depth and spatial coverage, our platform advances whole-brain transcriptomics, enabling 3D neuron typing capability critical for deciphering brain-wide cellular architectures.

## 1. Introduction

Neuron typing is critical for advancing our understanding of brain function and diseases [1-3]. Extensive studies have revealed that individual neurons or specific neuronal subtypes project across multiple brain regions, uncovering intricate interregional connectivity patterns and functional architectures [4-9]. Furthermore, population-level dynamics often originate from single-cell heterogeneity, where variations in neuronal firing patterns, synaptic plasticity, and gene expression collectively shape the emergent functions of neural networks [10].

Current approaches for neuron typing include morphological typing, electrophysiological typing, and molecular typing, each revealing distinct aspects of neuronal diversity and complexity [11-16]. Among these, molecular typing offers unique advantages as certain molecular characteristics remain stable throughout a cell’s lifespan, serving as reliable genetic markers. More specifically, this approach enables the construction of molecular atlases by delineating the correlation between molecular signatures and spatial distributions, thereby providing a framework for elucidating distinct neural circuit connectivity and functional dynamics [1-3]. Notably, molecular typing enables precise mapping of cellular distribution and functional characteristics across different brain regions and hierarchical levels in intact brain tissue sections. Therefore, molecular typing elucidates cross-hierarchical and trans-regional neural network architectures, offering substantial insights into understanding brain functions and neural circuit mechanisms [17-19].

Fluorescence in situ hybridization (FISH) constitutes a cornerstone technology for molecular typing. This technology achieves precise spatial mapping of different kinds of RNA molecules within tissue sections, enabling identification of distinct neuronal subtypes through their unique gene expression patterns [17, 18, 20-24]. Recent technical innovations, including but not limited to MERFISH, osmFISH, ClampFISH and STARmap, have expanded imaging capabilities across large tissue volumes [19, 24-28]. However, most of current implementations remain constrained to thin tissue sections (10–20 μm in thickness), limiting molecular typing to two-dimensional cellular distributions that incompletely reconstruct three-dimensional neural architectures and impede analysis of intercellular interactions [18, 21, 24].

Recently, researchers have been trying to develop FISH-based molecular typing methods for thicker tissue sections. For example, Fang et al combined MERFISH with spinning-disk confocal microscopy to achieve three-dimensional imaging of specific brain regions (100–200 μm thickness) with 100 ms exposure time per optical section [29]. However, despite improved acquisition speed and reduced photobleaching in the spinning-disk confocal microscopy, the inherently low transmission efficiency of emission fluorescence through the disk apertures results in insufficient signal-to-noise ratio (SNR). This limitation often necessitates prolonged exposure times, paradoxically diminishing overall imaging throughput [30]. On the other hand, Wang et al combined light-sheet microscopy with tissue expansion, and developed a new molecule typing technique, called EASI-FISH, which achieved RNA profiling in small-volume tissue blocks (300 μm thickness, 0.8 mm × 0.8 mm area) [31]. However, the dependence of EASI-FISH on expansion microscopy techniques may lead to excessive volumetric expansion of intact thick brain sections, potentially causing non-uniform probe penetration [21, 31, 32] and increasing detection volume which substantially elevates both workload and complexity of spatial registration.

After considering the above information, we realized that various intrinsic limitations restrict current molecular typing methods to either analyzing localized (millimeter-scale area) thick-tissue samples or processing complete (centimeter-scale area) thin-section specimens, thus creating persistent technical challenges for achieving high-resolution three-dimensional mapping of RNA distributions in intact and thick brain sections.

To address this challenge, here we propose a new FISH-based neuron typing method for high-precision RNA imaging in intact brain sections with 50 μm thickness and centimeter-scale lateral coverage. This method, called List-FISH, combines tilted light-sheet illumination with linear scanning to enable imaging of intact brain sections while maintaining throughput [9]. By replacing conventional thin light-sheets with thick light-sheets, uniform illumination is achieved in specimens with tens-of-micron thickness [33]. Notably, while traditional light-sheet microscopy links axial resolution to light-sheet thickness, List-FISH enhances axial resolution through astigmatic point spread function (PSF) encoding and spatiotemporal decoding (localization) of FISH signals within light-sheets of several microns in thickness, achieving sub-micrometer resolution in 3D [34, 35]. Furthermore, as data acquisition within each imaging column is inherently continuous, intra-column registration becomes obsolete, necessitating only inter-column alignment. This fundamental optimization significantly reduces registration complexity and enhances data processing efficiency.

Using experiments in both fluorescent bead phantoms and murine brain sections, we demonstrated that List-FISH could enable high resolution and high throughput RNA imaging in intact 50 μm-thick brain sections. We performed comparison between the reconstructed spatial data and the Allen Mouse Brain Atlas to further demonstrate essential consistency in global gene expression patterns, thereby validating the method’s accuracy in resolving spatial RNA distributions within intact thick brain sections. This paper provides an efficient and reliable solution for studying RNA distributions in intact thick brain sections, while establishing a significant foundation for exploring gene functional relationships in complex intact thick-tissue specimens.

## 2. Method

### 2.1 Optical path settings

The optical system comprises three functionally distinct modules: (i) light sources, (ii) wide-field imaging, and (iii) light-sheet imaging (Fig. 1). The light sources consists of four lasers with wavelengths of 405/470/532/640 nm (Laserwave, LWVL405-200mW, LWBL470-2W, LWGL532-4W, LWRL640-2W). After collimation by lens groups L1 and L2, the laser beams are combined via dichroic mirrors DM1-DM3 (Chroma, ZT532rdc-UF1, ZT473rdcxt-UF1, ZT405rdc-UF1). A mirror motorized on a flip mount FM (Thorlabs, MFF101) is employed to switch the optical path between wide-field and light-sheet imaging modalities. When the mount is detached, the laser beam is first expanded by lenses L3 and L4, and shaped into a 45°oblique light sheet by cylindrical lenses CL1-CL3, and focused onto the sample through a glass tapered light pipe GC for thick-tissue illumination. Fluorescence from the sample is collected via objective Obj1 (Nikon, CFI Apo NIR 40X W) that is positioned at 90° relative to the illumination axis.

**Fig. 1.**
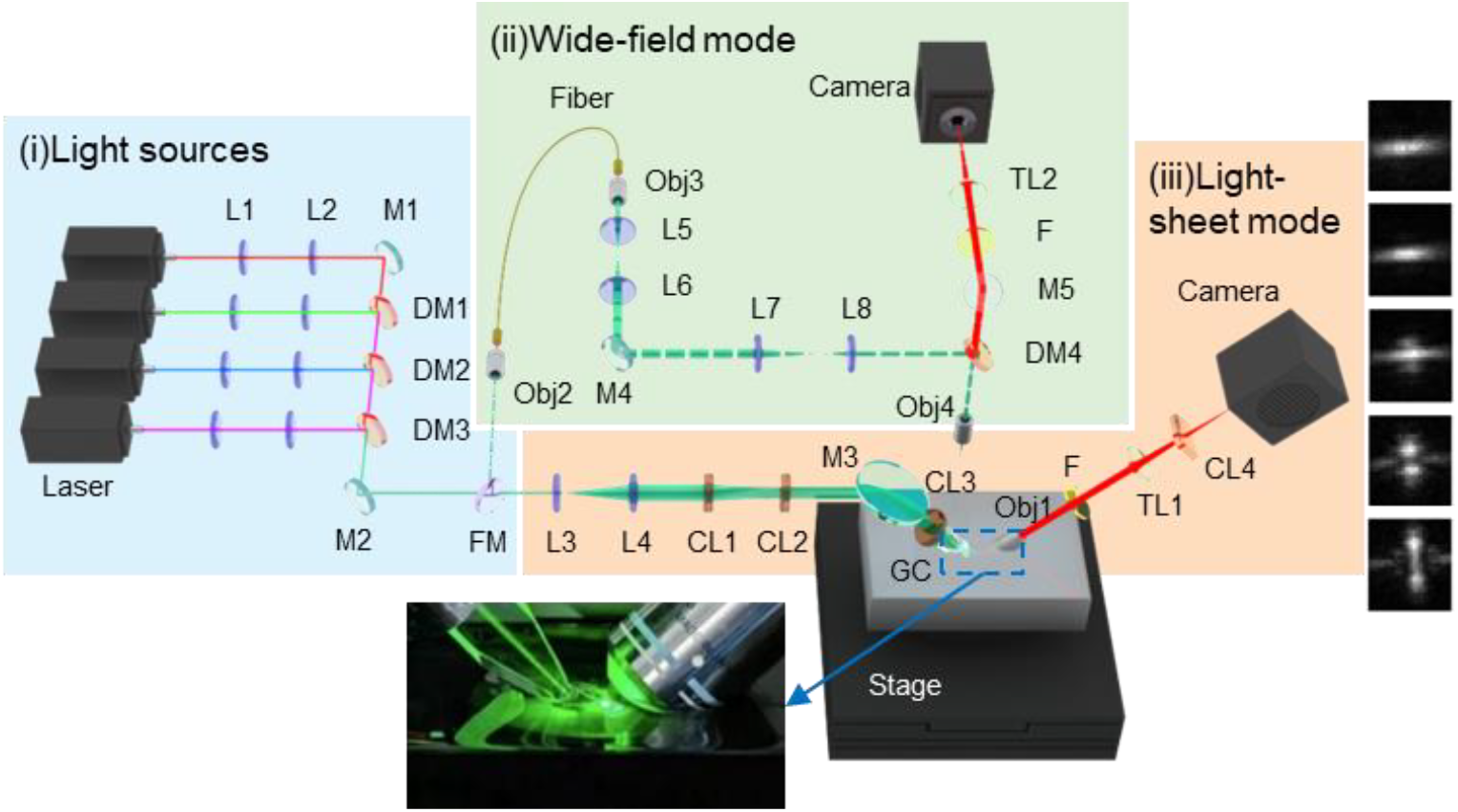
Schematic of the optical path.

Fluorescence signals pass through filter F1 (Chroma, ZET405/473/532/640) to block laser, then are modulated by tube lens TL1 and cylindrical lens CL4 to generate astigmatic point spread functions (PSF) on a sCMOS camera (Hamamatsu Photonics, C13440-20CU). In the wide-field unit, FM is used to redirect the laser beams, and a 200 μm-square fiber is used to couple the beams through objectives Obj2 and Obj3 (Olympus, PLN10X). The beams are expanded by lenses L5-L8, reflected by dichroic mirror DM4 (Chroma, ZT405/488/532/640rpc-XT-UF1), and projected as a square illumination field via objective Obj4 (Olympus, UMPLFLN20XW). Fluorescence signals collected by Obj4 pass through filter F2 (identical to F1) and are imaged via tube lens TL2 on another sCMOS camera (Hamamatsu Photonics, C13440-20CU). A high-precision XYZ stage (AEROTECH, ANT130XY-160 & AVL125) is used to enable sample positioning, synchronized with image acquisition and storage via a workstation.

### 2.2 Spatiotemporal localization algorithm

The spatial information of FISH spots within thick light-sheets is determined by a home-built spatiotemporal localization algorithm called LF-LOC, which is implemented through the following workflow:

***Step 1***: 2D localization. Raw images collected from the camera in the light-sheet unit are transferred into the LF-LOC workflow (Fig. 2) to perform 2D localization, which includes image preprocessing, ROI identification, and 2D localization using MLE-LM (Maximum Likelihood Estimation via Levenberg Marquardt method) [35], so that the lateral coordinates (x, y) and morphological parameters (σ_x_, σ_y_) of FISH spots can be simultaneously determined. This process generates a 2D localization table (x, y, σ_x_, σ_y_,…), where the lateral positions and morphological descriptors serve as critical inputs for subsequent axial localization.

**Fig. 2.**
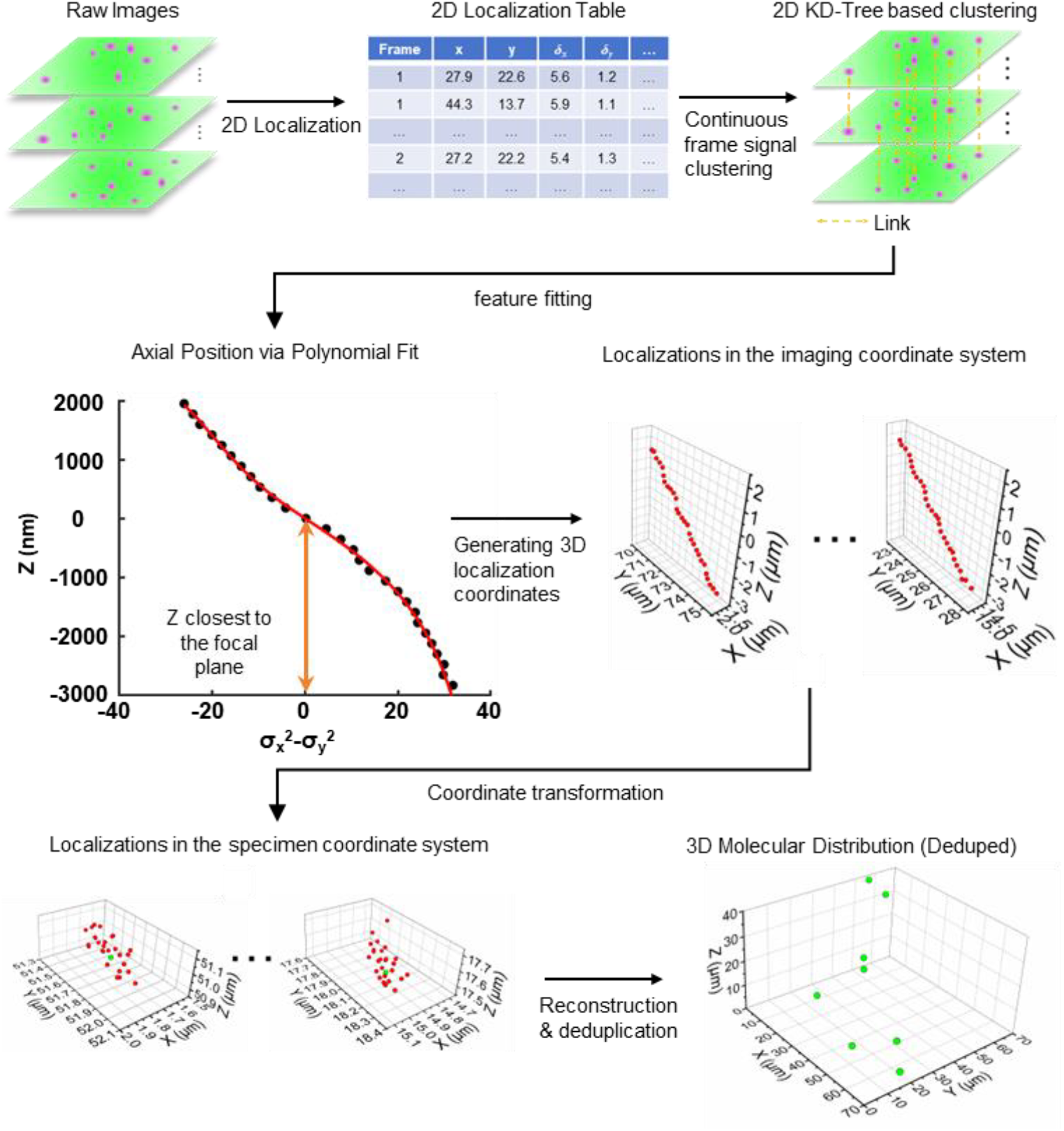
Flowchart of the spatiotemporal localization algorithm.

***Step 2***: Continuous frame signal clustering. Building upon the 2D molecular coordinates obtained in Step 1, this study employs a KD-tree-based spatiotemporal clustering method to link and integrate signals of the same FISH spot across consecutive frames, processing their spatiotemporally extended data. While precise Z-axis coordinates remain unresolved at this stage, the method successfully constructs consolidated fluorescence signal clusters, providing a preprocessed foundation for subsequent 3D localization.

***Step 3***: Feature fitting. For each FISH spot, the PSF ellipticity metric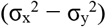 derived from spatiotemporally upsampled data is subjected to sixth-order polynomial fitting against the axial positions (Z-axis) retrieved from the motorized stage.

***Step 4:*** Generating 3D localization coordinates. The fitted axial positions are intercept-subtracted to calibrate the origin offset, generating a series of 3D coordinates within the light-sheet spatial domain.

***Step 5:*** Coordinate transformation. The 3D localization coordinates generated in Step 4 are actually within the imaging coordinate system (that is, the light-sheet imaging coordinate), and should undergo affine transformation to the specimen coordinate system, so that all localizations are precisely aligned with the stage positions recorded during acquisition. This transformation yields a specimen-referenced 3D localization table that maintains spatial fidelity to the biological sample.

***Step 6***: Reconstruction & deduplication. The frame index corresponding to the minimal absolute value of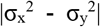 is selected for each FISH spot, discarding coordinate points associated with other feature values. By integrating the synchronized stage coordinates recorded during acquisition of the selected frames, high-precision 3D localization in the specimen coordinate system is achieved. This curated dataset subsequently enables visual 3D reconstruction of RNA distributions at subcellular resolution. The detailed information is in Supplement 1.

### 2.3 Imaging fluorescent beads

We used fluorescent bead samples to optimize the imaging parameters and evaluate the imaging performance of our List-FISH method. We conducted experiments using a four-step protocol.

***Step 1***: Sample preparation. A 5% (w/v) agarose solution was prepared by heating distilled water to dissolve agarose powder. This solution was then used to dilute 100 nm fluorescent microspheres (Invitrogen, F8800) to a final concentration of 1 × 10^−8^ (one hundred millionth of the original concentration). After a complete dispersion of the microspheres, 50 μl of the mixture was uniformly coated onto a glass slide and allowed to settle for 1 minute. This coating process was repeated eight times to incrementally increase the sample thickness. The sample was subsequently dried in an oven at 60°C for 12 hours to fully cure the agarose matrix. After post-curing, structural fiducial markers were created by selectively removing portions of the gel-embedded microspheres using a sharp tool, ensuring stable spatial references for subsequent imaging validation.

***Step 2***: Imaging region selection. First, the sample was secured in the specimen chamber, followed by the injection of 2× SSC solution as the imaging buffer. Subsequently, the high-precision motorized stage mentioned in Section 2.1 was driven to position the sample beneath the objective lens of the wide-field mode. The Z-axis position was adjusted to achieve optimal imaging clarity. A tile-scan imaging protocol was then executed with a 500 μm lateral step size (X-Y plane) to scan the entire signal-containing region, and the acquired sequential images were stitched into a composite panoramic image. Based on this composite image, the target region of interest (ROI) was identified, and its spatial boundary coordinates were mapped within the stage coordinate system. These coordinates were subsequently transformed into the light-sheet imaging coordinate system using the pre-calibrated spatial offset between the wide-field and light-sheet objectives, thereby precisely defining the volumetric imaging boundaries for high-resolution light-sheet imaging.

***Step 3***: Light-sheet imaging. First, the continuous scanning length for each light-sheet along the X-axis was set in the control software, and the motorized stage was actuated to move at a constant velocity. During stage motion, the camera acquired images at an interval of 266 nm (X-axis sampling interval) and an exposure time of 2.5 ms per frame. Subsequently, the number of scan columns along the Y-axis was set, with a step size of 300 μm between each column, to achieve full coverage of the detection area. During image data acquisition, the image acquisition system simultaneously recorded the motorized stage coordinates corresponding to each image frame, ensuring precise spatial correspondence.

***Step 4***: Confocal microscopy imaging. The sample was subjected to 3D confocal imaging using a commercial microscope (Zeiss, LSM710) equipped with a 40× oil-immersion objective, with Z-axis optical sectioning performed at 0.5 μm step size to maintain spatial registration with the pre-scanned region.

### 2.4 Imaging mouse brain slices

The mouse brain slice samples used in the experiments were obtained from Shulaibao (Wuhan, China) Biotechnology Co., Ltd., and processed through standardized protocols to guarantee experimental reproducibility.

The RNA staining was designed and improved from the our previous work [36]. Briefly, the brain slices were fixed with 4% PFA at room temperature (RT) for 15 min, digested with 5 μg/ml Proteinase K at 37 for 20 min, transferred into permeabilization buffer (2×SSC, 2% v/v sodium dodecyl sulfate (SDS), 0.5% Triton-X-100) for 10 min, and then stored in Methanol at-20°C overnight. The next day, the brain slices incubated in 1× hybridization buffer (2× SSC, 35% formamide, 1% Triton-X-100, 0.5% SDS, 20mM RVC, 50 nM per oligo) at 37 °C overnight. The brain slices were ligated by T4 DNA Ligation mixture (0.2 U/μl T4 DNA Ligase supplemented with 0.2U/μl RRI) at 4°C for 12h, moved to RT for 3 hours, and amplified by Phi29 DNA polymerase mixture (0.2 U/μl Phi29 DNA polymerase, 250 μM dNTP, 0.2 U/μl RRI and 10nM 5-(3-aminoally1)-dUTP) at 4°C for 6h, moved to 30°C for 3 hours with gentle shaking. Then, the brain slices were modified by Acrylic acid NHS ester at RT for 4h and embedded with Acrylamide hydrogel overnight. Finally, the tissue gel was digested with Proteinase K to reduce the background fluorescence.

For RNA detection, we employed a multi-round imaging approach targeting the following genes: *Cck, Cnr1, Npy, Cux2, Reln, Calb1, Lgi2, Penk, Rorb, Slc17a7, Crym*, and *Rprm1*. For each round, 3 read-out probes were hybrid with 2×SSC, 10% v/v ethylene carbonate incubated at RT for 30 min. After imaging, samples were incubated with buffer with 10% TCEP (Tris(2-carboxyethyl) phosphine hydrochloride) and washed with 2×SSC to remove the fluorescent signal that coupled with disulfide bond in the fluorescent oligos.

After each imaging round, the imaging buffer was drained from the specimen chamber, and 500 μL TCEP solution was dispensed onto the sample and allowed to stand for 15 minutes. Residual TCEP was then drained, and 50 mL 2× SSC was introduced into the chamber for two sequential 10-minute washes. Subsequently, a hybridization mixture containing 470 μL hybridization buffer (2× SSC, 10% vinyl carbonate) and three RNA-FISH read-out probes (10 μL each, 10 μmol stock) for the next imaging targets was applied to the sample and incubated for 1 hour. After draining residual probe-containing hybridization buffer, three 5-minute washes with 500 μL fresh hybridization buffer were performed. Finally, 50 mL 2× SSC was introduced for a 2-minute incubation before drainage, completing the probe removal and re-staining cycle.Following the completion of the elution and re-staining procedures, the imaging process and subsequent elution/re-staining steps were repeated until all pre-defined imaging rounds were fully completed.

## 3. Results and discussions

### 3.1 The imaging depth of List-FISH

While conventional light-sheet microscopy enhances axial resolution through light-sheet thickness minimization, a strategy inherently limiting maximum sample thickness [37], List-FISH employs an intentionally thickened light-sheet (Fig. 1) to maintain uniform illumination across extended tissue volumes. This design fundamentally trades axial resolution for enhanced illumination homogeneity. To resolve this limitation, we introduced a cylindrical lens anterior to the sCMOS camera, encoding fluorescence PSF variations across the illumination volume via depth-dependent astigmatism. Quantitative analysis of astigmatic PSF morphology enables submicron axial resolution recovery, effectively compensating the thick light-sheet’s inherent resolution penalty. This strategy draws inspiration from single-molecule localization microscopy principles [34, 35], now adapted for volumetric imaging in scattering tissues.

Notably, List-FISH employs a detection objective oriented orthogonally to the light-sheet plane. This configuration enables each acquired frame to resolve FISH signals within the 45°-tilted volume illuminated by the light sheet (Fig. 3(a)). This geometric relationship necessitates coordinate system transformation between light-sheet imaging coordinate system and specimen coordinate system for accurate thickness quantification. Fluorescent bead samples were employed for systematic imaging depth validation through single-frame acquisitions. Quantitative characterization in Fig. 3(a) reveals a longitudinal signal distribution width of 156.0 μm. Subsequent geometric calibration yielded a sample thickness of 109.6 μm.

**Fig. 3.**
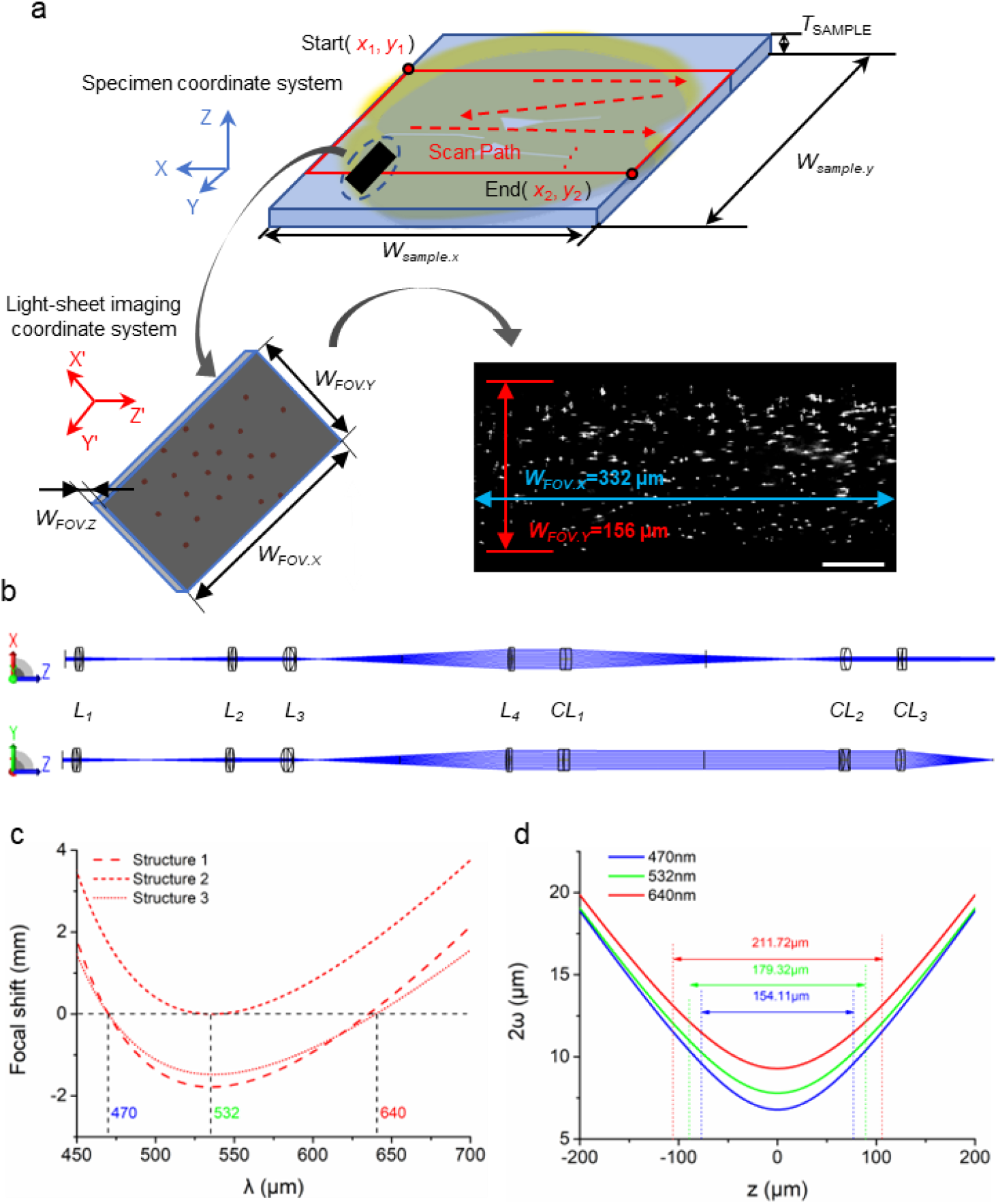
Optical sectioning methodology and illumination characterization in List-FISH. (a) In the schematic of the continuous axial scanning strategy, the Rayleigh range (black rectangle) defines the effective imaging depth. The imaged volume under light-sheet illumination allows sample thickness estimation by registering the imaging and specimen coordinate systems. For example, a single-frame raw fluorescence bead image shows a signal FWHM of 156.0 μm, corresponding to a sample thickness of 109.6 μm. Scale bar: 50 μm. (b) Zemax-optimized sequential optical path with beam transformation stages: collimation (L1-L2), beam expansion (L3-L4), and light-sheet shaping (CL1-CL3). (c) Multi-configuration simulation demonstrating chromatic focal shift elimination through air gap parameterization. (d) Light-sheet thickness (2ω(z)) versus propagation distance, showing wavelength-dependent divergence rates (λ=470/532/640 nm).

To systematically characterize light-sheet behaviors, we implemented a linear sequential optical model in Zemax (Fig. 3(b)), where ray widths were proportionally scaled for illustrative purposes only to visualize lens-induced beam convergence/divergence. The air gap between L1 and L2 was parameterized as a variable. Configurations with primary wavelengths of 470 nm, 532 nm, and 640 nm were established via the multi-configuration function, achieving focal position alignment across wavelengths and consequent elimination of chromatic focal displacement (Fig. 3(c)).

The simulated beam widths (2ω(z)) for the three wavelengths are plotted as functions of propagation distance z in Fig. 3(d) using the equation below:

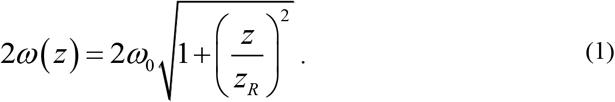

Quantitative characterization demonstrates that 470 nm excitation achieves the minimal depth of focus (154.1), with geometrically calibrated effective sample thickness measuring 107.9 μm. Progressive expansion to 179.3 μm and 211.7 μm occurs at 532 nm and 640 nm excitations respectively, exhibiting nearly linear correlation with wavelength while maintaining effective thickness >100 μm across all configurations. Combined experimental and theoretical analyses confirm that the system’s volumetric imaging capability exceeds the 100 μm threshold while maintaining a dimensional deviation of less than 2% from theoretical predictions.

### 3.2 The 3D resolution of List-FISH

As detailed in Section 3.1, spatiotemporal evolution of signal morphology is systematically recorded through sequential raw image acquisition. These time-resolved intensity profiles enable precise three-dimensional localization via the LF-LOC algorithm, achieving axial resolution enhancement beyond conventional light-sheet microscopy’s inherent thickness limitation. This computational approach establishes a robust framework for high-resolution volumetric imaging of thick biological specimens (> 100 μm), where traditional optical sectioning methods prove inadequate.

We quantified the 3D spatial resolution of List-FISH through systematic characterization of fluorescent bead localization precision. We also benchmarked the axial localization performance between our LF-LOC algorithm and a conventional calibration curve-based axial analytical method [34].

First, we processed 3,439 sets of raw data (corresponding to 3,439 FISH spots, and each spots generate tens of fluorescent signals) using MLE-LM [35] to localize the 2D planar coordinates and extract morphological features of fluorescent signals in each frame. Subsequently, we applied both the calibration curve-based axial analytical method and the LF-LOC algorithm to resolve the centroids of these localized fluorescent signals. We further performed clustering on the localization table containing all localization information for each FISH spots and aligned the centroids of these clusters in 3D space. Finally, we characterized their distributions using histogram statistics and Gaussian fitting along the x, y, and z axes. Note that the standard deviation (σ) derived from the fitting can be used to calculate the FWHM (full width at half maximum) resolution of the imaging technique [34, 38].

As shown in Fig. 4(a-b), compared to the 3D point cloud distribution generated by the calibration curve-based method, the results from the LF-LOC algorithm exhibit significantly narrower clustering along the Z-axis, demonstrating superior axial localization precision. Fig. 4(c-d) displays the histogram statistics and Gaussian fitting results for the X- and Y-axis directions, revealing a FWHM resolution of 275 nm at X axis and 350 nm at Y axis, respectively, from the List-FISH method. Fig. 4(e-f) quantitatively compares the Z-axis localization precision between the two algorithms. The calibration curve-based method achieved an axial FWHM resolution of 1664 nm, whereas the LF-LOC algorithm substantially improved this precision to 559 nm, unequivocally valSubsequently, the same region of bead idating the algorithm’s superior 3D localization performance for List-FISH data.

**Fig. 4.**
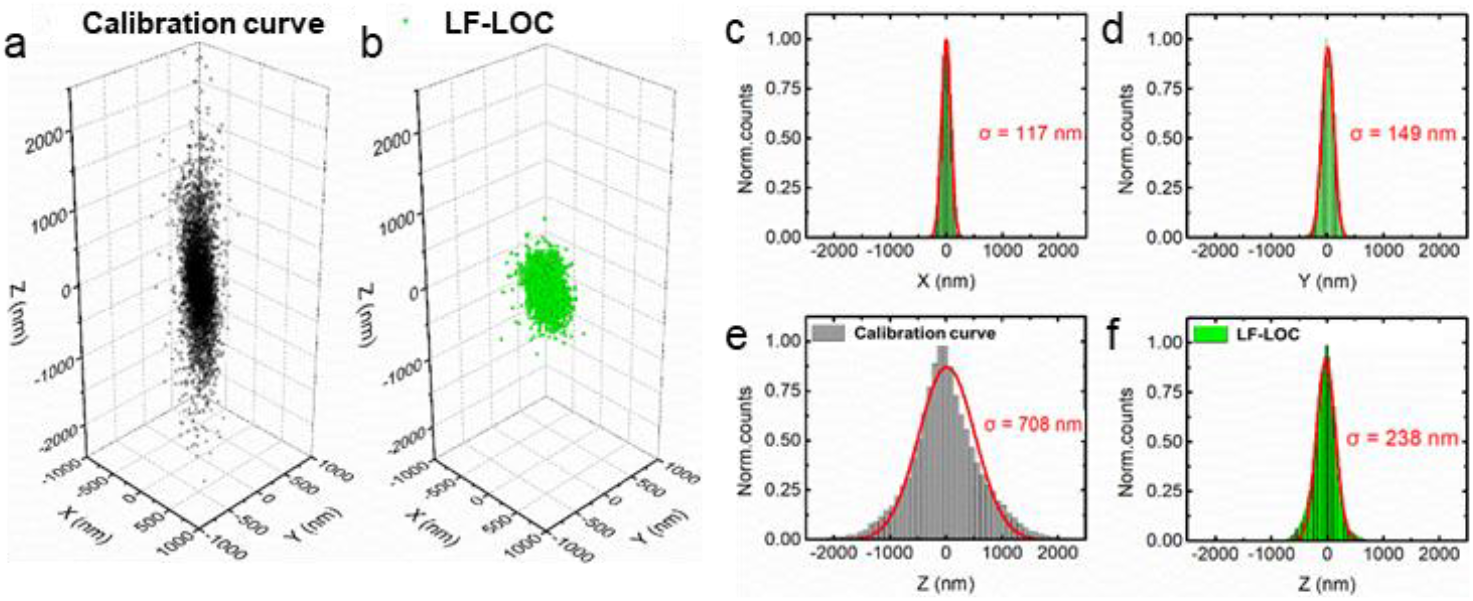
Localization distribution of fluorescent signals from a total of 3,439 FISH spots. (a) The distribution of localizations from calibration curve-based axial analytical method. (b) The distribution of localizations from the LF-LOC algorithm developed in this study. The same localization algorithm (MLE-LM) was used to localize the XY positions of the fluorescent signals presented in (a) and (b). (c-d) Histogram statistics of the localizations in (b) using Gaussian fitting in x/y directions, yielding σ values of 117 nm and 149 nm, respectively. (e) Z-direction histogram analysis with Gaussian fitting for the calibration curve-based axial analytical method (σ = 708 nm). (f) Z-direction histogram analysis with Gaussian fitting for the LF-LOC algorithm (σ=238 nm). Note that FWHM = 2.35σ [34, 38].

### 3.3 Comparing the imaging performance of List-FISH with confocal microscopy

We first imaged fluorescent bead samples using the List-FISH method and then performed confocal microscopy imaging with the same sample region. We utilized confocal microscopy images as the ground-truth benchmark, which were employed to evaluate the imaging performance of List-FISH with the Jaccard coefficient [38]. This approach also enabled us to assess the detection accuracy of the LF-LOC algorithm and further demonstrate its feasibility and advantages within the List-FISH framework.

The raw data in List-FISH imaging of fluorescent bead samples were processed using the LF-LOC algorithm to reconstruct 3D spatial coordinates of beads through coordinate transformation. The reconstructed data were then axially projected onto a 2D plane and rendered via 2D Gaussian fitting (Fig. 5(a)). Subsequently, the same region of bead sample was imaged using confocal microscopy, and the axial maximum intensity projection (MIP) of the confocal results was displayed in Fig. 5(b). The images in Fig. 5(a) and Fig. 5(b) were fused by color channels to produce the merged image in Fig. 5(c). Note that Figs. 5(a–c) show maximum intensity projections in the XY plane. Fig. 5(d) highlights a detailed region from Fig. 5(c). In Fig. 5(d), 83 molecules (see orange boxes) localized by LF-LOC were successfully matched with confocal counterparts, while 2 mismatched molecules (cyan boxes) and 1 undetected molecule (purple boxes) were identified.

**Fig. 5.**
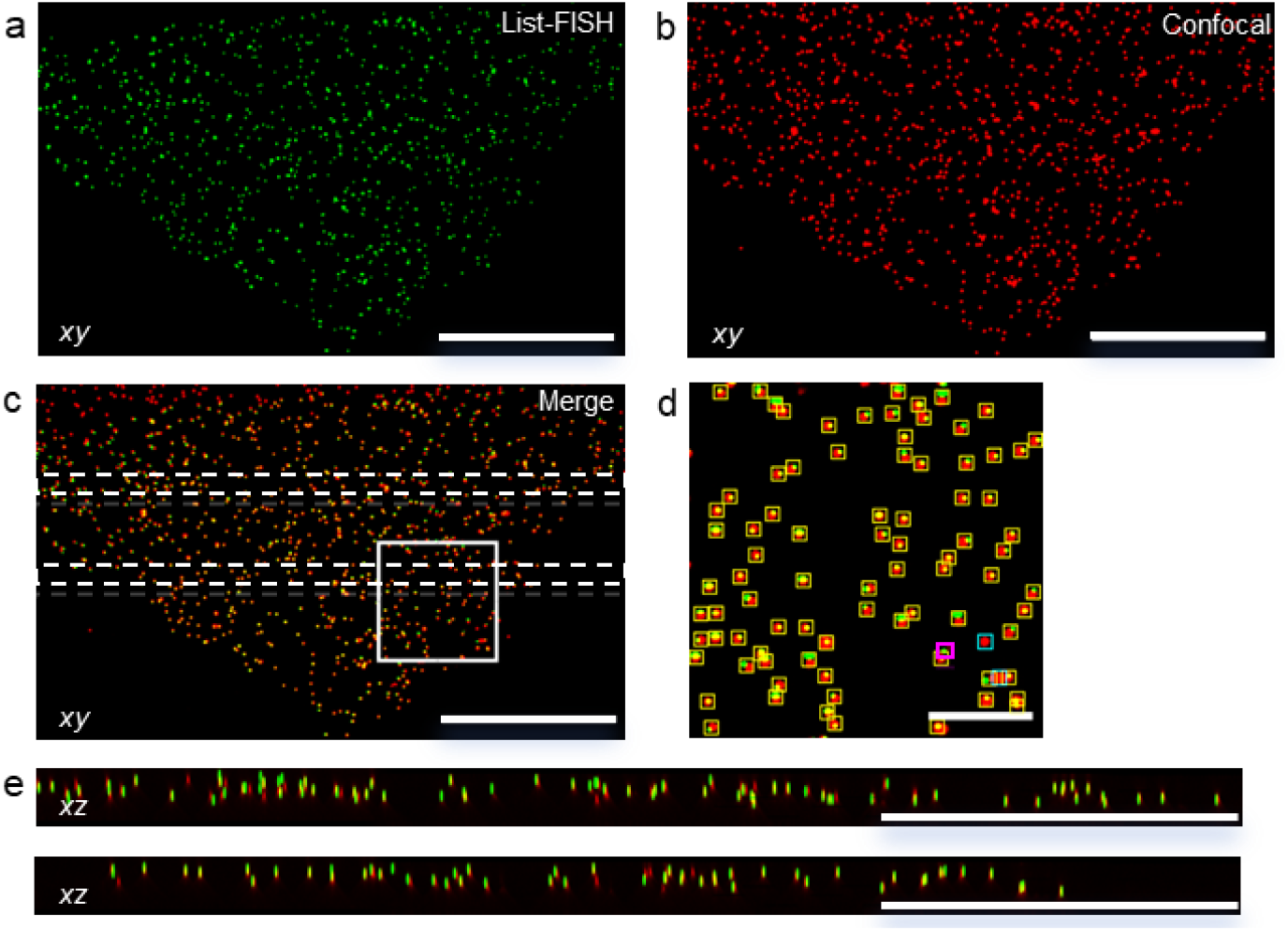
Comparative analysis of List-FISH reconstructions and confocal imaging results. (a-b) List-FISH localization reconstruction (a) and confocal imaging result (b) of the identical model specimen. (c) Merged image of (a) and (b). (d) Magnified view of the white-boxed region in (c), with orange boxes indicating true positives (TP, n=83), cyan boxes denoting false negatives (FN, n=2), and purple boxes marking false positives (FP, n=1). (e) Magnified view of the central region between the white dashed lines region in (c), Scale bar: 100 μm.

The XZ view of the central region between the white dashed lines in Fig. 5c is shown in Fig. 5e, which also presents axial cross-sections of the merged data from both imaging systems. In this axial section, 113 true positives (TP), 10 false positives (FP), and 3 false negatives (FN) were identified. The resulting Jaccard coefficient for molecular identification reached 97% (XY) and 90% (XZ)—a high benchmark in molecular localization research[38]. Considering the inherent coordinate system difference between the two imaging systems and the nonlinear change of samples with time, the molecular overlap rate in the merged image may decrease. These results in Fig. 5 could validate LF-LOC’s high accuracy in fluorescent spot detection and its feasibility and advantages for List-FISH imaging.

### 3.4 Mouse brain imaging

To confirm the high-resolution RNA imaging performance of the system in thick tissue samples, List-FISH experiments were conducted in intact mouse brain slices. Although the previous model sample imaging experiments (see Section 3.1) have confirmed that the system supports the imaging of tissue sections exceeding 100 μm in thickness. The imaging thickness is also limited by fluorescence labeling method: the signal consistency and reconstruction accuracy would be a major concern in thick-tissue fluorescence labeling, because uniform probe penetration and efficient hybridization become increasingly constrained in thicker tissues due to limited permeability. Considering the trade-off between imaging depth and FISH signal quality, a slice thickness of 50 μm was selected to ensure homogeneous probe distribution, stable fluorescence signal, and reliable localization results.

In a pilot study of whole-brain List-FISH imaging experiment performed on a 50 μm-thick section, three gene targets—Hap1, Sst, and Mbp—were simultaneously detected. From the resulting 3D dataset, a 10 μm-thick axial sub-volume was selected for high-resolution localization analysis. This sub-volume was then projected along the Z-axis to generate a corresponding 2D gene expression map (Fig. 6(a)). To further assess the accuracy of the spatial expression profiles, the resulting distribution was compared with reference data from the Allen Brain Atlas. The experimental results (Fig. 6(b)) exhibited a high degree of spatial concordance, validating the system’s capability for RNA localization and spatial mapping in intact brain tissue.

**Fig. 6.**
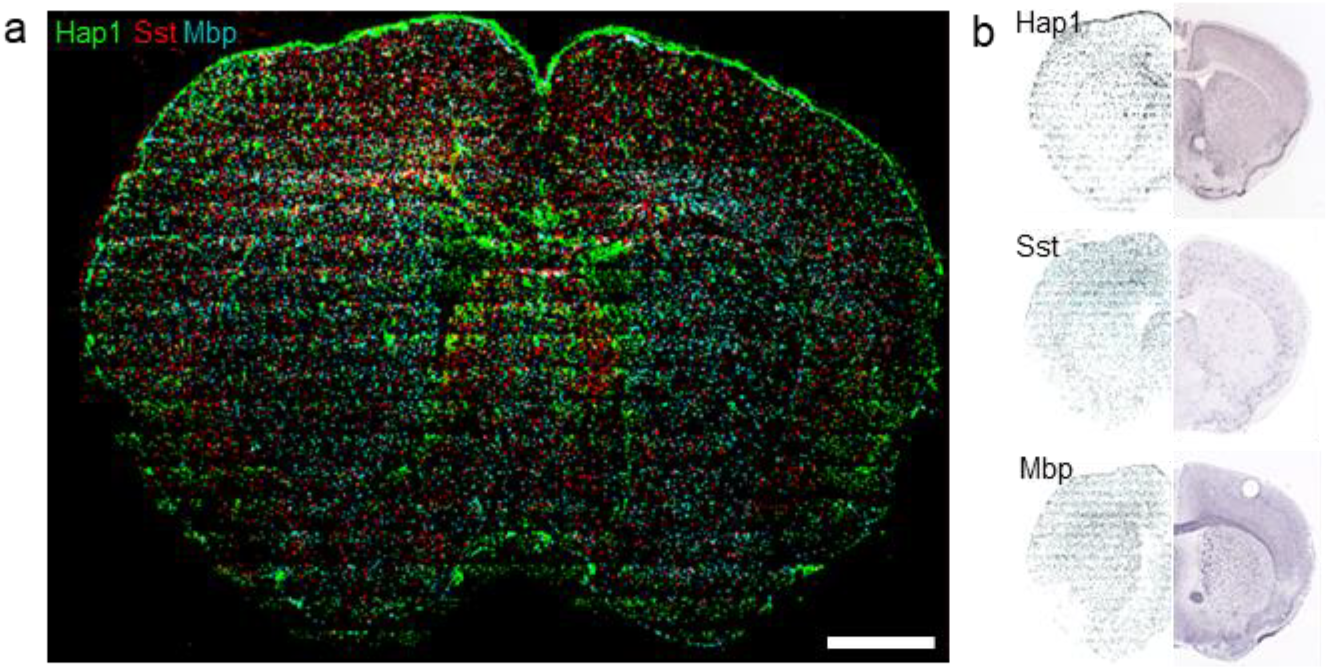
Spatial mapping of representative RNA genes in 50 μm-thick intact brain tissue. (a) List-FISH mapping of Hap1, Sst, and Mbp. A 10 μm axial segment was projected along the Z-axis to generate the 2D expression map. Scale bar: 1 mm. (b) Comparison of List-FISH results with the Allen Brain Atlas show strong agreement in spatial gene expression patterns.

It is noteworthy that some stripe-like artifacts are visible in the reconstructed images. These artifacts are primarily attributed to intensity inhomogeneities of the Gaussian excitation beam used in the laser scanning system, as well as imperfections in post-processing correction algorithms. Further optimization of the imaging and computational pipeline is currently underway.

To further investigate the 3D RNA imaging performance of List-FISH, a pilot study was performed on a 50 μm-thick mouse brain tissue section, focusing on a selected cortical region. Through three rounds of List-FISH imaging, spatial expression patterns of seven genes—Rorb, Penk, Lgi2, Calb1, Reln, Cux2, and Npy—were detected (Fig. 7(a)). Cell segmentation was conducted using the open-source Cellpose algorithm [39]. As shown in Fig. 7(b), the spatial distributions of gene expression across different axial depths of the 50 μm-thick sample are visualized. The first panel presents a full-depth projection in which all axial layers were collapsed into a single 2D image, resulting in the superimposition of cellular structures from various depths and the loss of laminar information. The subsequent five panels show sectional images taken at 10 μm intervals, revealing the layered architecture along the Z-axis. These images demonstrate clear variations in cellular density and distribution across different depths, with RNA signals predominantly clustered along the inner boundaries of individual cells. Enabled by cell segmentation, this clustering pattern becomes more clearly observable, supporting the biological plausibility and subcellular accuracy of the localization results. Additionally, RNA localizations exhibit pronounced spatial heterogeneity between neighboring cells, highlighting the orderly spatial arrangement of distinct cell types within the tissue volume.

**Fig. 7.**
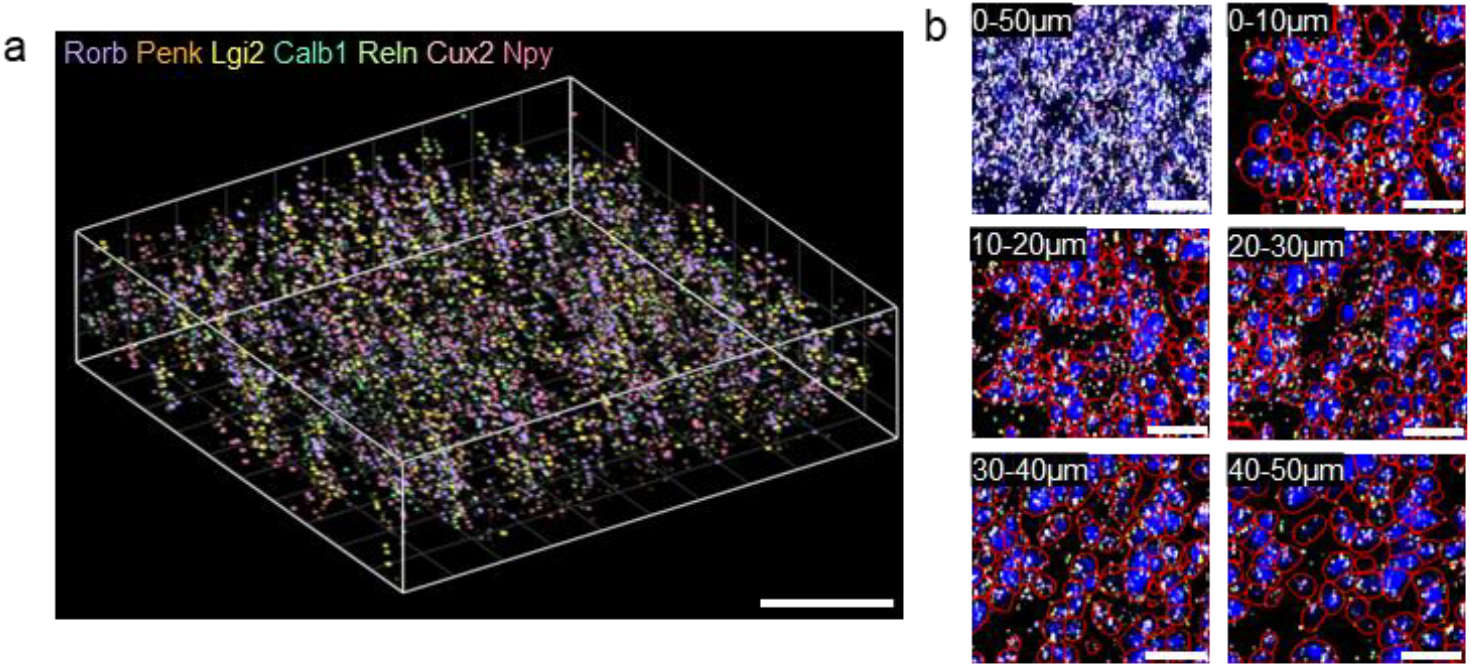
High-resolution 3D RNA mapping in a 50 μm-thick cortical region. (a) 3D reconstruction of a cortical region showing spatial localization of seven genes (Rorb, Penk, Lgi2, Calb1, Reln, Cux2, Npy) detected over three List-FISH rounds. Scale bar: 50 μm. (b) Axial projections of the same region. The first panel shows a full-depth projection, while the remaining panels display 10 μm-thick sections at different depths. Cell segmentation using Cellpose reveals RNA clustering near the inner cell boundaries (red lines). Scale bar: 50 μm.

## 4. Conclusion

In this paper, we developed a FISH-based molecular typing method termed List-FISH. This method employs tilted thick light-sheet illumination to achieve uniform illumination across thick specimens, while astigmatic PSF and a LF-LOC algorithm ensure high-resolution RNA localization capability. Compared to existing RNA imaging methods applicable to intact brain slices, List-FISH supports imaging of tissue sections of up to 100 μm thickness, enabling the acquisition of 3D spatial distributions of RNAs within intact brain tissues. This advancement provides a powerful tool for studying cellular spatial organization and RNA distribution in complex tissues, offering novel insights into the relationship between spatial architecture and functional dynamics of cells in neuroscience research.

We validated the system’s imaging capability for samples of >100 μm in thickness using dried agarose gel samples embedded with 100 nm fluorescent beads. The lateral (X/Y) and axial (Z) localization errors were controlled to be within 117 nm, 149 nm, and 238 nm, respectively, demonstrating high localization accuracy. Compared to confocal microscopy results (serving as ground truth), the system achieved a Jaccard coefficient of 97% (X/Y) and 90% (X/Z) for molecular identification—a high benchmark in molecular localization research. List-FISH enables accurate and high-resolution RNA localization in intact brain tissue sections up to 50 μm thick, supporting both large-scale two-dimensional spatial profiling and detailed three-dimensional reconstruction. In a pilot single-round imaging, gene expression patterns across entire brain sections closely reflected known anatomical structures, validating the system’s performance in preserving spatial fidelity. Multi-round hybridization further expanded detection capability without compromising image quality or localization accuracy.

A pilot study on high-resolution 3D RNA imaging of selected cortical regions revealed biologically meaningful spatial features, including perinuclear RNA clustering and laminar neuronal organization along the axial dimension. These observations highlight structured spatial heterogeneity in transcript distribution and reflect underlying subcellular positioning and tissue-level architecture.

Together, these findings demonstrate that List-FISH is well-suited for scalable and multiplexed spatial transcriptomics, providing a robust platform for high-fidelity molecular mapping in complex tissue environments.

## Acknowledgements

We thank all the other members of Digital Theranostics and Optical Microscopy (DigiTOM) group from Hainan University for their technical support.

## Data availability

Codes and data underlying the results presented in this paper are not publicly available at this time but may be obtained from the authors upon reasonable request.

## Supplemental document

See Supplement 1 for supporting content.

